# A new hypothesis for type 1 diabetes risk: The at-risk allele at rs3842753 associates with increased beta cell *INS* mRNA in a meta-analysis of single cell RNA sequencing data

**DOI:** 10.1101/2020.12.06.413971

**Authors:** Su Wang, Stephane Flibotte, Joan Camunas-Soler, Patrick E. MacDonald, James D. Johnson

**Affiliations:** Diabetes Research Group, Life Sciences Institute, Department of Cellular and Physiological Sciences & Department of Surgery, University of British Columbia, Vancouver, Canada, V6T 1Z3; UBC-LSI Bioinformatics Core Facility, University of British Columbia, Vancouver, British Columbia, Canada; Department of Bioengineering, Stanford University, Stanford, CA 94305, USA; Department of Pharmacology and Alberta Diabetes Institute, University of Alberta, Edmonton, Canada

**Keywords:** type 1 diabetes, genetics, insulin expression, VNTR, single cell RNAseq

## Abstract

Type 1 diabetes is characterized by the autoimmune destruction of insulin secreting β cells. Genetic variations upstream at the insulin (*INS*) locus contribute to ~10% of type 1 diabetes heritable risk. Multiple studies showed an association between rs3842753 C/C genotype and type 1 diabetes susceptibility, but the molecular mechanisms remain unclear. To date, no large-scale studies have looked at the effect of genetic variation at rs3842753 on *INS* mRNA at the single cell level. We aligned all human islet single cell RNA sequencing datasets available to us in 2020 to the reference genome GRCh38.98 and genotyped rs3842753, integrating 2315 β cells and 1223 β-like cells from 13 A/A protected donors, 23 A/C heterozygous donors, and 35 C/C at-risk donors, including adults without diabetes and with type 2 diabetes. *INS* expression mean and variance were significantly higher in single β cells from females compared with males. Comparing across β cells and β-like cells, we found that rs3842753 C containing cells (either homozygous or heterozygous) had the highest *INS* expression. We also found that β cells with the rs3842753 C allele had significantly higher ER stress marker gene expression compared to the A/A homozygous genotype. These findings support the emerging concept that inherited risk of type 1 diabetes may be associated with inborn, persistent elevated insulin production which may lead to β cell ER stress and fragility.

## INTRODUCTION

Pancreatic β cells are the body’s primary source of insulin, a hormone that promotes glucose uptake from the bloodstream into tissues [1]. Type 1 diabetes constitutes ~10% of diabetes cases and is caused by the autoimmune destruction of a substantial proportion of β cells [1]. Type 1 diabetes is highly heritable with a twin concordance between 30% and 70% [2]. The HLA locus is the major genetic determinant of type 1 diabetes risk (OR = 0.02 to 11), but other genes play a significant role as well [3]. Gene mapping and GWAS studies identified the human insulin (*INS*) locus as the 2^nd^ most important contributor of type 1 diabetes genetic risk (OR = 2.38, ~10% of heritable risk) [4; 5]. The region of the *INS* locus implicated in type 1 diabetes susceptibility has three genetic variants in near-perfect linkage disequilibrium with each other [4–6]. Variants associated with increased type 1 diabetes risk are: a shortening of the repetitive region 400bp upstream of *INS* (class III *INS-*VNTR > class I *INS*-VNTR), a single nucleotide polymorphism (SNP) in the *INS* 5’ intronic region (rs689 T>A), and a SNP in the *INS* 3’UTR (rs3842753 A>C) [4–8]. Often, the SNP alleles are examined as surrogates for *INS*-VNTR length [6; 8–10]. The rs3842753 C allele has been consistently correlated with increased type 1 diabetes susceptibility [5; 6; 8].

Despite genetic studies linking the rs3842753 C allele to increased type 1 diabetes susceptibility, there have only been three studies examining the effect of these genetic variants on *INS* expression in the pancreas [6; 9; 10]. The studies found an allele specific association between rs3842753 C allele and increased *INS* expression in heterozygous adult and fetal whole pancreas tissue, but were confounded by a lack of cell type specificity. Moreover, the sample sizes were small (n = 3, 1, and 10) [6; 9; 10]. These findings lend support to the β cell ER stress model of type 1 diabetes pathogenesis. It has been hypothesized that increased insulin demand could generate large proportions of misfolded or unfolded proinsulin, leading to ER stress [11]. Under glucose stimulation [12] and even basal conditions [13], insulin production is a significant source of ER stress. If the allele-specific increase in *INS* expression is translated into increased insulin production, there could be increased ER stress activated unfolded protein response leading to increased generation of unique antigen that present exclusively under specific conditions (neoautoantigens) [14]. Neoautoantigens derived from *INS* have been shown to stimulate T-cell proliferation and cytokine production, though the role of these neoautoantigens on type 1 diabetes autoimmunity remain unclear [15].

Recently, multiple data sets of single cell RNA sequencing (scRNAseq) from human pancreatic islet cell have been produced [16–23]. Studies have found distinct subpopulations of β cells and β-like cells, with varying conclusions for differences in *INS* expression, confirming heterogeneity as a core feature of islet cell biology [18; 23]. Studies have also found transcriptomic differences in β cells from donors without diabetes compared to donors with type 2 diabetes [16–18], although these have not always been reproducible between studies [24] (likely due to the complex type 2 diabetes etiology and limited sample size). We reasoned that integrating all available scRNAseq data from human pancreatic islets into one dataset and conducting analyses with a greater sample size could resolve true differences.

In this study, we examined the effect of rs3842753 genotype, sex, and type 2 diabetes status on *INS* expression independently and in combination in single pancreatic β cells using an integrated dataset containing all available scRNAseq data. Our findings are consistent with the novel hypothesis that elevated β cell insulin production contributes to inherited type 1 diabetes risk by increasing β cell stress.

## METHODS

### Data inclusion

We conducted all scRNAseq alignments and initial dataset integration analysis using the Cedar Compute Canada cloud compute resource (www.computecanada.ca). All subsequent analyses were done in RStudio version 3.6.0. We included all published pancreatic islet scRNAseq datasets obtained using Smart-seq2 or Smart-seq methods for library preparation due to their higher sequencing depth for accurate SNP genotyping. We also included the Human Pancreas Analysis Program (HPAP) data available at the time of our study [22]. We obtained the published datasets through the Gene Expression Omnibus (GEO) or the European Bioinformatics Institute (EMBL-EBI), and the HPAP datasets directly from investigators (Supplemental Table 1). All dataset metadata contain donor age, sex, and diabetes status. The read length ranges from 43bp to 100bp and median read depth per cell ranges from 0.75 to 4.4 million reads. In total, we used data from 71 donors, split into 48 donors without diabetes and 23 donors with type 2 diabetes. At the time of our analysis, single cell data was only available from 1 adult donor and 1 child donor with type 1 diabetes, precluding their use in our analysis which requires multiple donors of each genotype. We also expect that gene expression analysis of β cell from donors with type 1 diabetes would be severely confounded by the stresses associated with the disease.

### Read alignment and genotyping

Single cell RNAseq reads were aligned to human reference genome GRCh38.98 using STAR version 2.7.1a [25] with gene read counts obtained using --quantMode GeneCounts. Read count files were aggregated into study specific read count matrices using a custom code. Each cell was genotyped at SNP rs3842753 (chr 11:2159830), using Samtools mpileup version 1.9 [26]. Pancreatic β cells (indicated by total read count (DP4) at rs3842753 > 100) were used for donor genotype determination. Reference allele percentage was calculated as 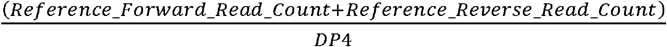. The alternate allele percentage was calculated as 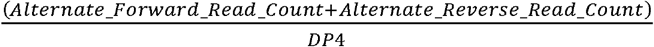. Homozygosity was determined as reference or alternate allele percent > 80%, and heterozygosity otherwise. Donor genotype is determined as the majority of the individual’s β cells’ genotype. Less than 5% of individual β cells’ genotype are non-concordant with donor genotype. Genotype non-concordant cells are included in downstream analysis and re-labeled with donor genotype.

### Dataset filtering and cell type analysis

Initial filtering removed cells with abnormally high total RNA counts and genes expressed (i.e. doublet cells) and cells with low number of expressed genes and/or high mitochondrial gene percent (low viability), through visual inspection of distribution graphs for each dataset. The upper and lower limits were specific to each dataset to account for library preparation and sequencing protocol differences. The highest mitochondrial gene percentage was set to 25%. Datasets were integrated into a single dataset in Seurat version 3.1 [27] using SCTransform [28] and clustered in UMAP space using default settings. Top ten differentially expressed genes in each cluster (compared to all other cells) was used to determine cluster cell type identify. As well, the location of pancreatic hormone genes (*GCG, INS, PPY* and *SST*) expression was used to determine Alpha (α), β, PP, and delta cell clusters, respectively. Enrichr (gene enrichment analysis) [29] was used when cluster identity could not be determined using previous methods.

### Gene expression analysis

ER stress marker genes expression was normalized to all genes expressed in cell according to default Seurat settings. *INS* gene expression was normalized against select housekeeping genes rather than against all expressed genes in a cell due to the overwhelming amount of *INS* expression in pancreatic β cells (50% of all read counts) and to prevent self-normalization. Housekeeping genes for *INS* expression normalization were selected from a list of established human housekeeping gene (*ACTB, GAPDH, PGK1, PPIA, RPLP0, B2M, SDHA, TFRC, GUSB, HMBS, HPRT1, TBP*) for low inter-dataset variance and sufficient expression (relative to all genes in the cell). *INS* read counts in β cells and β-like cells were normalized against the sum of select housekeeping genes read counts, scaled to a total of 10000, and natural log transformed. All cells with zero values for *INS* read counts or select house-keeping genes read counts were excluded from the downstream analysis. We analyzed β cells and β-like cells separately. We checked for the expression of other genes in the rs3842753 linkage dis-equilibrium block (IGF2, TH), but did not find significant expression in single β cells, as expected.

### Statistical analyses

Shapiro-Wilk test was used to test for normality. Fligner-Killeen or Bartlett’s tests were used as a nonparametric or parametric test for difference in variance. Wilcox Rank Sum was used to test for differences in normalized *INS* expression between sexes or between disease statuses. Pairwise Kruskal-Wallis test was used to test for differences in normalized *INS* expression between genotypes, adjusting for multiple comparisons with Bonferroni correction. We did not study the effect of sex or diabetes status on normalized *INS* expression within genotype.

## RESULTS

### Dataset genotype summary

VNTR length cannot be determined directly with commonly used scRNAseq methods because reads are too short to span such large repetitive regions. Therefore, for each cell, rs384275 genotype was determined as a surrogate for *INS*-VNTR length and rs689 genotype due to the near perfect linkage disequilibrium between these gene variants in Caucasian populations. Also, there were no read alignments to the rs689 region. Within donors without diabetes, 9 donors had rs384275 A/A genotype, 12 had rs384275 A/C genotype, and 27 had rs384275 C/C genotype. Within donors with type 2 diabetes, 4 donors had rs384275 A/A genotype, 11 had rs384275 A/C genotype, and 8 had rs384275 C/C genotype (Supplemental Table 2).

### Cell type clustering and identification

We performed cluster analysis and labeling to identify specific cell types since our dataset included a mixture of cells from pancreatic islets. We found 14 distinct clusters within 13622 pancreatic islet cells (Figure 1A) and classified the clusters as α cells, β cells, PP cells, and delta cells based on expression of classical pancreatic hormone genes (*GCG, INS, PPY,* and *SST*) (Figure 1A, B, D). We also found endothelial cells, duct cells, and acinar cells based on *PECAM1, SPP1,* and *PRSS1* expression localization (Figure 1A, C, D). We identified mesenchyme and CD14+ monocytes using Enrichr based on top ten conserved genes per cluster compared to all other cells (Figure 1A, D). We identified a cluster of β-like cells that exhibited high *INS* expression compared to other clusters, but lacked the congregation in UMAP space. These cells may be considered poorly differentiated or dedifferentiated β cells, or they may be derived from other islet endocrine cells that have adopted *INS* expression. We used, separately, the β cell cluster (2315 cells) and the β-like cluster (1223 cells) for *INS* expression analysis.

**Figure 1:**
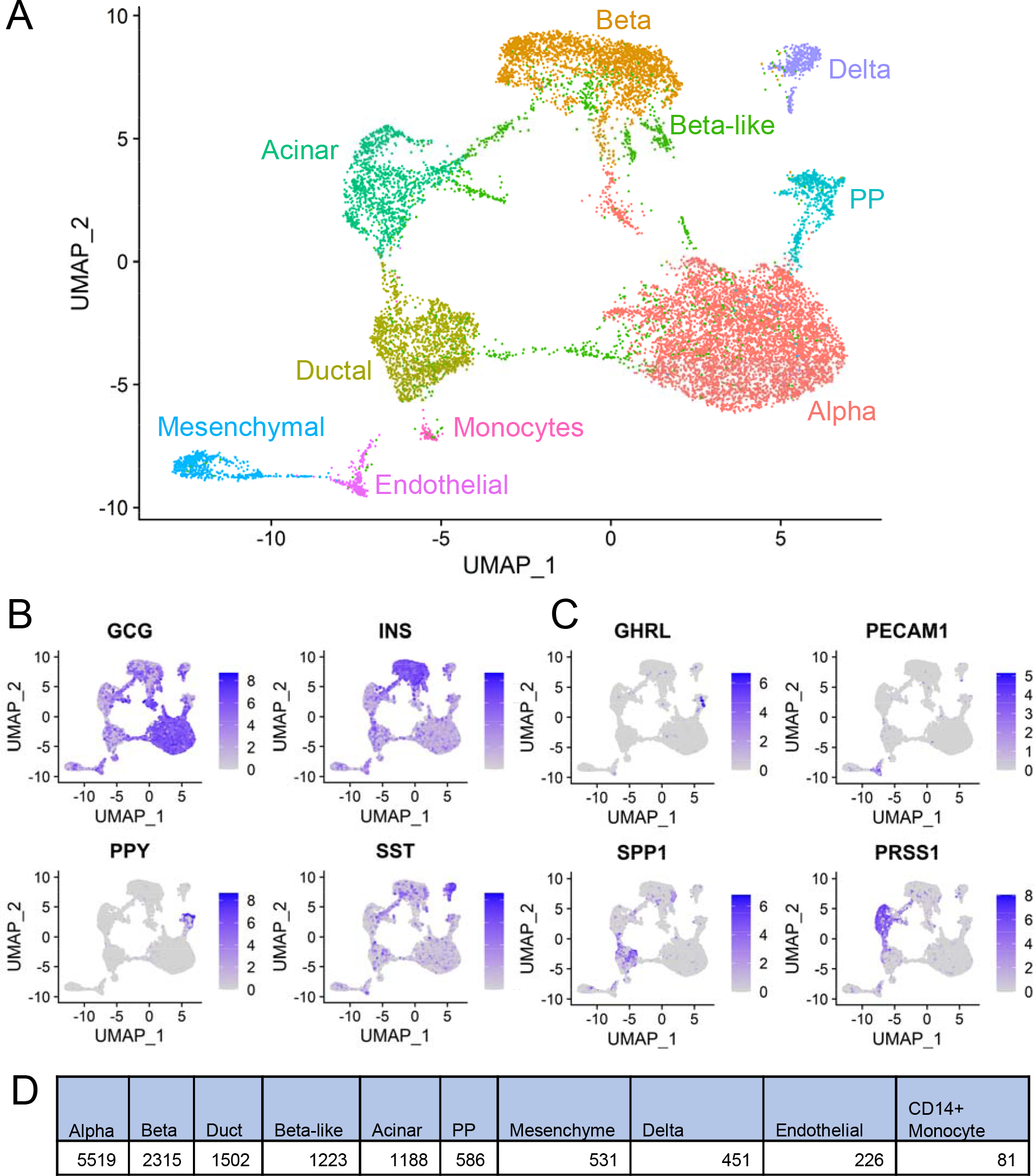
Cluster type clustering and identification of integrated dataset. UMAP projection of integrated dataset with cell type identity separated by colour (A). UMAP projection of integrated dataset with gene expression localization for pancreatic hormonal genes (B) and other cell type (C) markers. Each point represents one cell. (D) Table with each identified cluster cell number.

### Effects of female sex and type 2 diabetes status on INS mRNA in single β cells and /3-like cells

Sex and diabetes status have been shown to affect gene expression in pancreatic islets [16–18; 30]. We first examined the effect of sex on *INS* expression in β cells and β-like cells. In β cells, we found that cells from females and from donors with type 2 diabetes have increased normalized *INS* expression variance compared to cells from males and from donors without diabetes (Figure 2A, C). Interestingly, in β-like cells, we found the opposite association of an increase in normalized *INS* expression variance in cells from males and donors without diabetes compared to cells from females and from donors with type 2 diabetes (Figure 2B, D). In female donors, β cells, but not β-like cells, had increased levels of normalized *INS* expression compared to males (Figure 2A, B).

**Figure 2:**
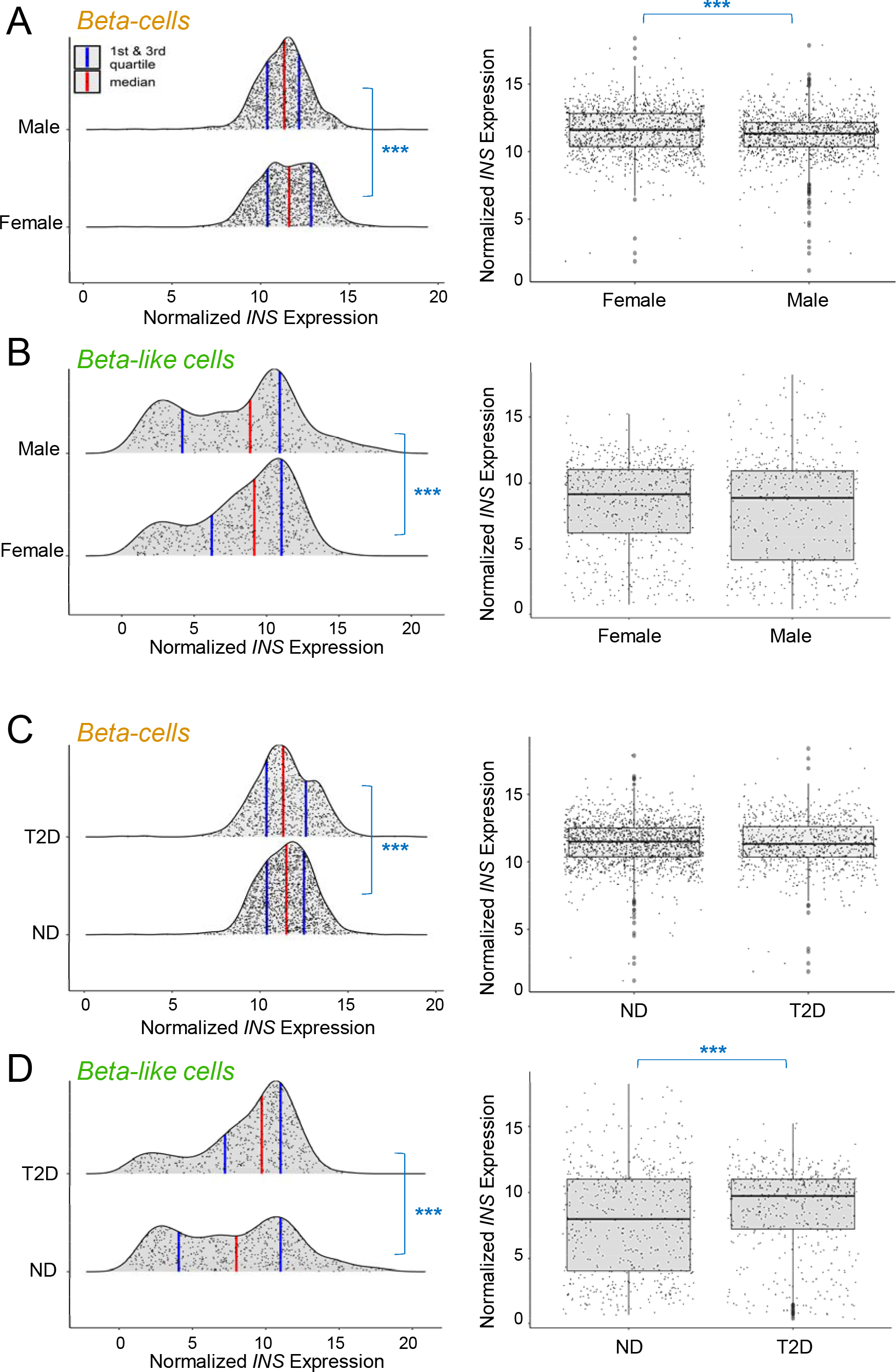
Normalized *INS* expression in be β cells and β-like cells by sex and disease status. Distribution and mean normalized *INS* expression separated by sex (in β cells: A, and in β-like cells: B) and disease status (in β cells: C, and in β cells: D). Red vertical lines represent median, blue vertical lines represent 1st and 3rd quartiles. Each point represents one cell. Bonferroni adjusted p < 0.001 = ***. ND = no diabetes, T2D = type 2 diabetes. Fligner-Killeen and Wilcox Rank Sum tests were used as nonparametric test for differences in variance and mean respectively.

Next, we examined whether *INS* expression in single β-cells was associated with type 2 diabetes. No significant difference in mean *INS* expression was observed between β cells from diabetic and nondiabetic donors (Figure 2C). In the β-like cells, normalized *INS* expression was increased in cells from donors with type 2 diabetes (Figure 2D). Changes in *INS* expression variance between sex and diabetes status provide additional evidence for heterogeneity in pancreatic β cell and β-like cell expression profile. Since *INS* expression level and variance changes between sex and diabetes status, it is important to account for these factors when studying the effect of rs3842753 genotype on *INS* expression.

### rs3842753 C allele associates with increased INS expression in single β cells and β-like cells

Previous studies used allele specific expression to show a correlation between the at-risk type 1 diabetes rs3842753 C allele and increased pancreatic *INS* expression in a maximum of 10 heterozygous whole pancreas tissues [6; 9; 10]. Here, we substantially increased the sample size (n = 71) and focused on *INS* expression specifically in β cells and β-like cells. To account for the overwhelming expression of *INS* in pancreatic β cells, and to avoid self-normalization to *INS*, we normalized *INS* expression to *B2M and RPLP0* expression, housekeeping genes we found to be both highly expressed and with a low interdataset variability in this dataset (Supplemental Figure 1). In β cells, donors with the rs3842753 A/C genotype had the highest level and variance of normalized *INS* expression, followed by C/C genotype, then A/A genotype (Figure 3A). Similarly, in β-like cells, cells with the at-risk C allele (A/C or C/C genotype) had the highest level of normalized *INS* expression (Figure 3B).

**Figure 3:**
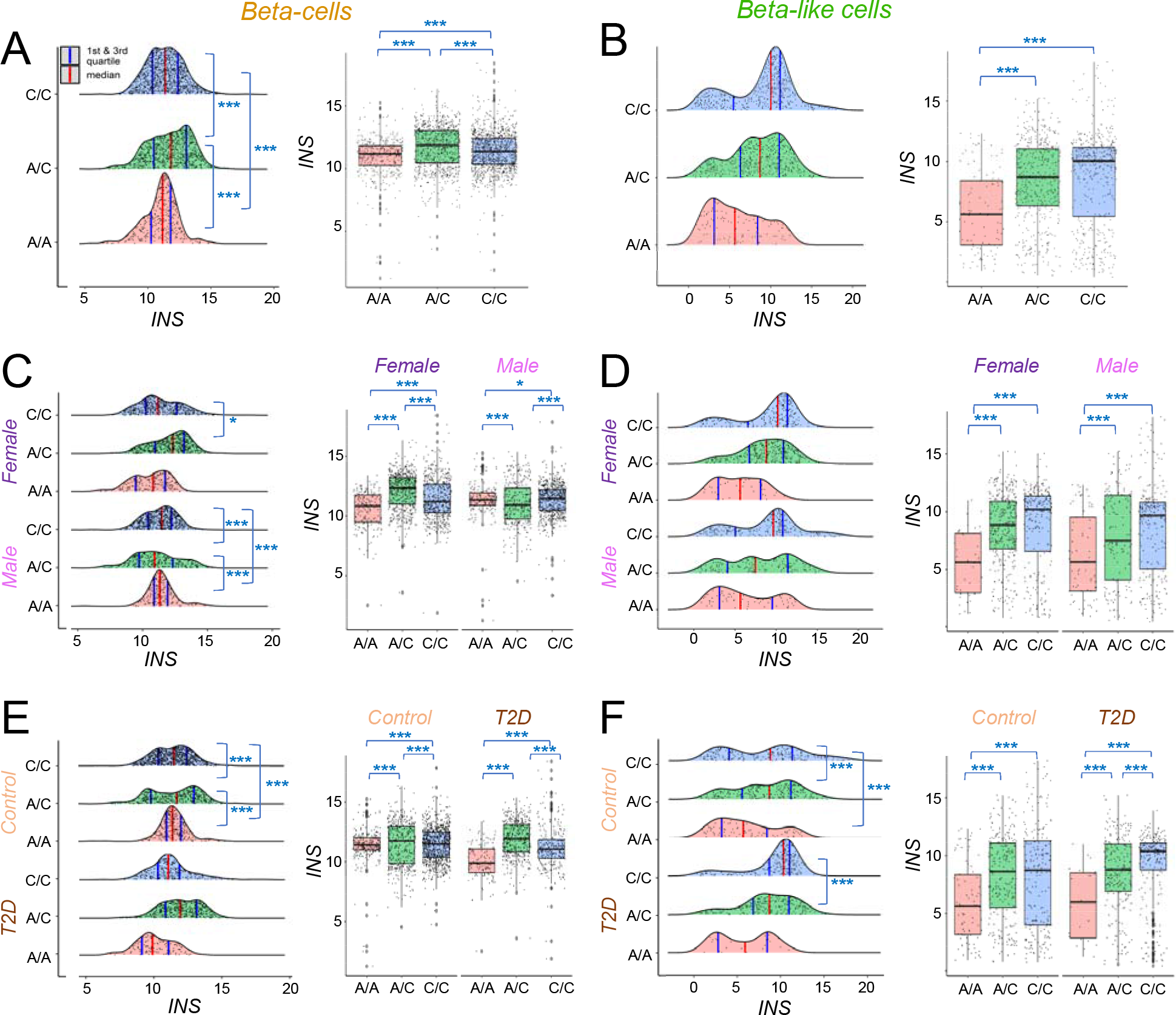
Normalized *INS* expression in β cells and β-like cells by genotype. Distribution and mean normalized *INS* expression separated by genotype (in β cells: A, and in β cell: B), genotype and sex (in β cells: C, and in β-like cells: D), and genotype and diabetes status (in β cells: E, and in β-like cells: F). Red vertical lines represent median, blue vertical lines represent 1st and 3rd quartiles. Each point represents one cell. Bonferroni adjusted P < 0.001 = ***. F = female, M = male, ND = no diabetes, D = type 2 diabetes. Graphs are coloured by genotype. Fligner-Killeen and pairwise Kruskal-Wallis tests were used as non-parametric test for differences in variance and mean respectively.

When examining the effect of rs3842753 genotype on *INS* expression within sex or diabetes status, β cells display the same trend of significantly increased variance of *INS* expression in donors with the heterozygous rs3842753 A/C genotype is present in cells from males and donors without diabetes (Figure 3 C, E). Additionally, cells from females and either diabetes status with the rs3842753 A/C genotype significantly expressed the highest level of *INS,* followed by C/C genotype, then A/A genotype (Figure 3 C, E). Regardless of sex or diabetes status, β-like cells also displayed increased *INS* expression associated with the rs3842753 C allele (either A/C or C/C genotype) (Figure 3D, F). Within either sex, there was no association between rs3842753 genotype and *INS* expression variance in β-like cells (Figure 3D). Similar to β cells, there was a significant increase in *INS* expression variance in β-like cells with rs3842753 C allele in donors with or without diabetes (Figure 3F). Together, these results indicate rs3842753 C allele affects *INS* expression, with the C allele associated with both increased *INS* expression variance and level in β cells and β-like cells. We attempted to examine the effects of rs3842753 on other genes within the linkage disequilibrium block but were unsuccessful due to the near zero expression of insulin like growth factor 2 and tyrosine hydroxylase.

### rs3842753 C allele associates with increased ER stress markers in single β cells and β-like cells

Insulin production has been shown to sustain chronic baseline ER stress [13]. Due to the increase in *INS* expression observed in cells with the at-risk rs3842753 C allele compared to the protective A allele, we were tested whether rs3842753 genotype also affected markers of ER stress. Indeed, the rs3842753 C allele was significantly associated with increased *ERN1* and *ATF6* expression in β cells (Figure 4A, B). There was no association between *ERN1* or *ATF6* expression and rs3842753 genotype in β-like cells (Figure 4C, D). These results indicate that rs3842753 C allele is associated with increased ER stress in ‘mature’ β cells.

**Figure 4:**
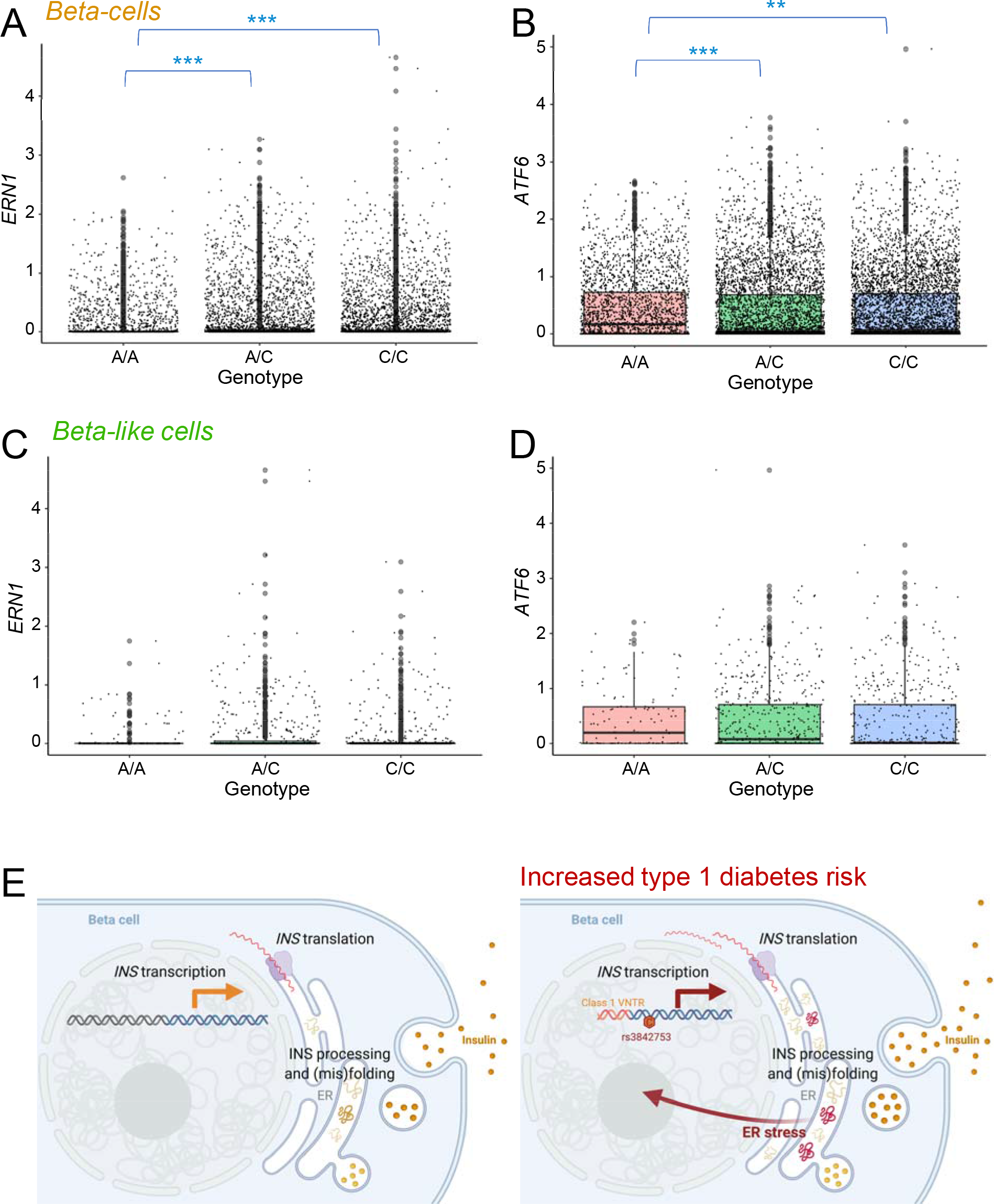
ER stress marked gene expression in β cells and β-like cells by genotype. Mean normalized ERN1 expression in β cells (A) and β-like cells (C), and ATF6 expression in β cells (B) and β-like cells (D), separated by genotype. Bonferroni adjusted P < 0.001 =***. Graphs are coloured by genotype. Pairwise Kruskal-Wallis tests was used as non-parametric test for differences in mean. (E) Working model–rs3842753 associates with increased INS mRNA levels, presumably resulting in increased INS translation and increased misfolded INS protein, leading to increased ER stress.

## DISCUSSION

The linked genetic variants at the *INS* locus: Class I INS-VNTR, rs689 A allele, and rs3842753 C allele, are the main variants associated with increased type 1 diabetes susceptibility [5; 6; 8]. Previously, only three small studies using whole pancreas tissues examined the effect of these variants on *INS* expression [6; 9; 10]. The studies accounted for inter-donor differences by examining allele specific *INS* expression in heterozygous donors. Our study is focused on the genetic factors that would precede type 1 diabetes. Beta-cells from type 1 diabetes donors would have been exposed to significant stress from type 1 diabetes and the results might not be representative of the pre-diabetic stage. Our study using 2315 single pancreatic β cells and 1223 β-like cells found that possession of a C allele at rs3842753 was associated with higher *INS* expression. These findings are consistent with the previous small studies using whole pancreas tissues that found that the C allele is associated with increased *INS* expression [6; 9; 10]. Sex distributions for previous studies were not reported. Our study included 71 adult donors (48 donors without diabetes and 23 donors with type 2 diabetes), which is a substantial improvement from previous studies with a maximum of ten fetal pancreas tissues.

Our study represents the largest compilation of human islets single-cell RNA sequencing data to date and it offers several insights into the biology of *INS* gene expression. As with the individual studies, our work re-enforces the substantial variability in insulin gene expression between cells. We also found that the heterozygous rs3842753 A/C genotype was associated with the highest variance of *INS* expression followed by A/A genotype, then by the C/C genotype in single pancreatic β cells. The increase in normalized *INS* expression variance in cells with A/C genotype could be due to the expression profile being influenced by both the rs3842753 A allele and C allele. This difference in variance supports the established model of heterogeneity of pancreatic β cells and perhaps the rs3842753 genotype plays a role in heterogeneity of *INS* expression [18; 23; 31]. Multiple other studies have found evidence of heterogeneity in β cells indicated by distinct subpopulations of β cells with varying *INS* expression [18; 23]. In β-like cells, variance of *INS* expression was less affected by the rs3842753 genotype. This is unsurprising due to the variety of cells that compose the β-like cells cluster, including polyhormonal cells and cells with gene expression signatures similar to α cells, β cells, ductal cells, and acinar cells (Figure 1A). Cell type variation in the β-like cells cluster was likely the primary factor affecting *INS* expression variance compared to the rs3842753 genotype. In other work from our lab, we have recently found that β cells can fluctuate between these states of high and higher insulin gene expression [31]. In the current study, we are unable determine whether comparatively high or low *INS* expression in single cells reflects dynamic or stable states.

Sex and diabetes status play an important role in insulin secretion and insulin resistance. We found that female donors had more pancreatic β cell *INS* expression than males. This is consistent with previous findings that males secrete less insulin after glucose stimulation than females, though insulin secretion under basal conditions is less extensively studied [32]. Interestingly, we found an increase in *INS* expression variance in β cells from females and donors with type 2 diabetes compared to males and donors without diabetes, respectively. This is consistent with the hypothesis that female pancreatic β cells need to have more plasticity in gene expression to accommodate the physiological demands of the potential for pregnancy and T2D β cells could have more variance due to the range in disease state and its effects on cell health and function. Conversely, β-like cells from males and donors without diabetes displayed greater variance in *INS* expression compared to cells from females and donors with type 2 diabetes, respectively.

Our study found that the rs3842753 C allele was associated with increased *INS* expression level in both β-like cells or β cells, though the specific genotype (A/C or C/C) associated with the highest *INS* expression level was different. In β-like cells, the rs3842753 C allele in either genotype (A/C or C/C) was associated with the highest *INS* expression level. Though not significant, except within β-like cells from donors with type 2 diabetes, β-like cells with the C/C homozygous genotype had the highest *INS* expression level followed by the A/C genotype, then the A/A genotype. Perhaps, the decreased overall *INS* expression level in β-like cells allowed for a great variance of *INS* expression, especially at higher *INS* expression levels, compared to ‘mature’ β cells. The effect of the rs3842753 genotype can be shown more clearly since the β-like cells are not expressing *INS* mRNA at their maximum capacity compared to ‘mature’ β cells.

In β cells, we made the perplexing finding that the highest *INS* mRNA expression was found in heterozygotes (A/C) donors. While we can only speculate, it is possible that such an overdominance mode of *INS* expression could be through a transcription factor competitive binding mechanism of action, proposed for other systems [33]. The rs3842753 C allele linked to shortening of *INS-*VNTR could increase accessibility of the chromatin by transcription factors at the *INS* loci and increase transcription of *INS*. At maximum *INS* expression capacity, transcription factor molecules in cells with the heterozygous genotype could exceed binding site availability, limiting competition. Cells with the homozygous allele have increased number of *INS* transcription factor binding sites, which theoretically could induce competition amongst binding sites for transcription factors (at *INS* loci or other locus) thus lowering transcription efficiency and decreasing *INS* expression. It would be interesting to conduct allele-specific *INS* expression analysis using SCALE or scBASE [34; 35]. However, we are unable to conduct those experiments using the current dataset due to the technical challenges of normalizing *INS* expression across multiple datasets with different sequencing protocols and donors.

Our study has several limitations. This includes sample size limitation in the areas of age range, ethnic demographic, and sample size, which means that the results should be cautiously extrapolated and multiple subgroup analysis is challenging. Most cases of type 1 diabetes onset between 10 to 14 years of age [1], but studied the *INS* expression from adults donors. Sufficient single islet cell data from children without diabetes or with early stages of type 1 diabetes is not available. Gene expression in β cells subjected to autoimmune attack would likely be difficult to interpret and confounded. The field of type 1 diabetes genetics in general is limited by an excessive focus on the Caucasian population [5; 6]. The near perfect linkage for the three genetic variants *INS*-VNTR, rs689, and rs3842753 has only been extensively studied in Caucasian populations with the correlation factor r^2^ unknown for other ethnicities [5]. Our dataset only included ethnicity metadata for 31 out of 71 donors (29 of Caucasian descent). Thus, our assumption that genotyping for SNP rs3842753 as a surrogate for *INS*-VNTR length could not be formally validated. Finally, despite compiling all available data at the time of our work, our study is still limited by the relatively small sample number (n = 71) compared to GWAS studies, though it is a major improvement compared to the previous studies (n = 1, 3, 10) [9]. Due to this limited sample size, we could not account for each of the large range of factors which could affect *INS* expression including: age, BMI, dietary differences, and ethnicity [1]. To address these limitations, we would need to increase the sample size, especially in younger age ranges.

We argue for an increase in studies using integrated scRNAseq datasets to take advantage of the massive amount of publicly available data. Our integrated dataset includes the most recent pancreatic islet scRNAseq data and will be available upon request. Diabetes status, sex, and age are included in the metadata allowing for investigations focusing on diabetes status specific transcriptome differences, sex specific transcriptome differences and age mediated transcriptome changes. Though Enge et al. (2017) have examined the transcriptional signatures of aging in human pancreas data with eight non-diabetic donors [19], it would be interesting to study the transcriptional signatures of aging between pancreatic islet cells from donors without diabetes compared to donors with type 2 diabetes. It would also be important to determine whether the changes in *INS* expression due to rs3842753 genotype we showed could be translated into changes in INS protein abundance. For this study, we could only obtain whole islet protein abundance data for the Camunas-Soler et al. (2019) and HPAP datasets [21; 22]. Though we observed the same trend in INS protein abundance as in *INS* expression (highest protein abundance in donors with rs3842753 C allele – homozygous or heterozygous), we could not conduct any statistical tests as there were only two donors with the A/A homozygous genotype (unreported). Future studies would benefit greatly with increased sample size and robust single-cell protein measurements, although it should also be noted that insulin protein content does not necessarily reflect transcription or translation rates.

What is the translational significance of these molecular genetic findings? Our results confirm the association between the rs3842753 C allele and increased *INS* expression, first proposed using whole pancreas from a small number of donors, at the single cell level using 2315 β cells and 1223 β-like cells from 71 donors. Our findings support the emerging concept that inherited risk of type 1 diabetes at SNP rs3842753 may be associated with elevated insulin production at the mRNA level and an increase in β cell ER stress. The insulin production-driven ER stress we have previously described [13] could make β cells, especially those near their maximal insulin synthesis and folding capacity, more fragile and sensitive to external stresses. Our data in β-like cells, which have lower absolute *INS* expression and do not display the same signs of ER stress, offers further support for that idea. Increased insulin production may also lead to errors in insulin mRNA translation, proinsulin protein processing and insulin peptide degradation, as well as other factors that could promote neoautoantigen generation and presentation, and the provocation of an autoimmune reaction (Figure 4 C) [11; 36]. Our results do not preclude a pathological role for VNTR-associated decrease in *INS* expression in the thymus, which has been proposed to impair central tolerance [37], although a dominant role for thymic negative selection in type 1 diabetes has been questioned [38]. Taken all together, paradoxically elevated pancreatic insulin production from birth could make some β cell more vulnerable to stress, against the background of HLA-driven autoimmunity. This hypothesis should be testable in animal models that have insulin production is specifically reduced in β cells [39], if they were crossed to a type 1 diabetes susceptibility background such NOD. If this hypothesis is further supported, it would provide even stronger rationale for approaches to reduce β cell stress in people who are at risk for type 1 diabetes.

## ACKNOWLEDGEMENTS

We thank Dr. Kaestner for providing access to the human Pancreas Analysis Program (HPAP) Database, consortia under Human Islet Research Network **(RRID:SCR_014393)**, funded by NIH grants **UC4-DK112217** and **UC4-DK112232**. We would also like to thank the donors and their families. This work was supported by a Canadian Institutes for Health Research grant to JDJ (PJT-152999). We would also like to thank the Anne and John Brown Fellowship in Diabetes and Obesity Related Research for funding this study.

## AUTHOR CONTRIBUTIONS

SW compiled and analyzed all the data, and wrote the manuscript. SF supported the data analysis. J C-S provided data and expertise. PEM provided data. JDJ conceived the study, edited the manuscript, and is the guarantor of this work.

