## Supplemental Table 1 for "A new hypothesis for type 1 diabetes risk: The at-risk allele at rs3842753 associates with increased beta cell *INS* mRNA in a meta-analysis of single cell RNA sequencing data"

**Supplemental Table 1: Dataset Sequencing and Accession Number.**

| Study | Date | Protocol | Sequencing System | Read Type | Read Length | Read Depth per Cell | Data Accession Location | Accession Method |
|---|---|---|---|---|---|---|---|---|
| Xin | 2016 | SmartSeq | Illumina HiSeq2500 | Single end | 75bp | 0.95 million | GSE81608 | GEO |
| Wang | 2016 | SmartSeq | Illumina HiSeq2500 | Single end | 100bp | 2.2 million | GSE83139 | GEO |
| Segerstolpe | 2016 | SmartSeq2 | Illumina HiSeq2000 | Single end | 43bp | 0.75 million | E-MTAB-5061 | EMBL-EBI |
| Camunas | 2020 | SmartSeq2 | NextSeq 500 or NovaSeq platform | Paired end | 75bp | 1 million | GSE124742 | GEO |
| Enge | 2017 | SmartSeq2 | Illumina NextSeq | Paired end | 75bp | 1 million | GSE81547 | GEO |
| HPAP | | | | | | | | |
| HPAP-1 | Date Accessed: September 2020 | SmartSeq | Illumina HiSeq2500 | Single end | 100bp | 4.4 million | https://hpap.pmacs.upenn.edu | provided by investigator |
| HPAP-3 | | | | | | | | |
| HPAP-4 | | | | | | | | |
| HPAP-6 | | | | | | | | |
| HPAP-7 | | | | | | | | |
| HPAP-8 | | | | | | | | |
| HPAP-10 | | | | | | | | |
| HPAP-12 | | | | | | | | |
| HPAP-13 | | | | | | | | |
