## Supplemental Table 2 for "A new hypothesis for type 1 diabetes risk: The at-risk allele at rs3842753 associates with increased beta cell *INS* mRNA in a meta-analysis of single cell RNA sequencing data"

**Supplemental Table 2: Donor number by sex and genotype for each dataset.**

| | | Non-Diabetic | | | | T2D | | | |
|---|---|---|---|---|---|---|---|---|---|
| **Study** | | **A/A** | **A/C** | **C/C** | **Total** | **A/A** | **A/C** | **C/C** | **Total** |
| Xin et al., 2016 (16) | **Total Donor #** | 3 | 4 | 5 | 12 | 1 | 3 | 2 | 6 |
| | **Male** | 3 | 1 | 3 | 7 | 0 | 2 | 1 | 3 |
| | **Female** | 0 | 3 | 2 | 5 | 1 | 1 | 1 | 3 |
| Wang et al., 2016 (17) | **Total Donor #** | 0 | 0 | 2 | 2 | 1 | 0 | 1 | 2 |
| | **Male** | 0 | 0 | 1 | 1 | 0 | 0 | 1 | 1 |
| | **Female** | 0 | 0 | 1 | 1 | 1 | 0 | 0 | 1 |
| Segerstolpe et al., 2016 (18) | **Total Donor #** | 3 | 0 | 3 | 6 | 1 | 2 | 1 | 4 |
| | **Male** | 3 | 0 | 2 | 5 | 0 | 1 | 1 | 2 |
| | **Female** | 0 | 0 | 1 | 1 | 1 | 1 | 0 | 2 |
| Camunas-Soler et al., 2020 (21) | **Total Donor #** | 1 | 4 | 13 | 18 | 1 | 4 | 2 | 7 |
| | **Male** | 1 | 2 | 6 | 9 | 0 | 2 | 1 | 3 |
| | **Female** | 0 | 2 | 7 | 9 | 1 | 2 | 1 | 4 |
| Enge et al., 2017 (19) | **Total Donor #** | 2 | 1 | 2 | 5 | Not Applicable | | | |
| | **Male** | 0 | 1 | 2 | 3 | | | | |
| | **Female** | 2 | 0 | 0 | 2 | | | | |
| HPAP (22) | **Total Donor #** | 0 | 3 | 2 | 5 | 0 | 2 | 2 | 4 |
| | **Male** | 0 | 1 | 1 | 2 | 0 | 0 | 1 | 1 |
| | **Female** | 0 | 2 | 1 | 3 | 0 | 2 | 1 | 3 |
| **Total Donor #** | | 9 | 12 | 27 | 48 | 4 | 11 | 8 | 23 |
| **Male** | | 7 | 5 | 15 | 27 | 0 | 5 | 5 | 10 |
| **Female** | | 2 | 7 | 12 | 21 | 4 | 6 | 3 | 13 |

Genotype at SNP rs3842753.
