## Supplemental Figure 1 for "A new hypothesis for type 1 diabetes risk: The at-risk allele at rs3842753 associates with increased beta cell *INS* mRNA in a meta-analysis of single cell RNA sequencing data"

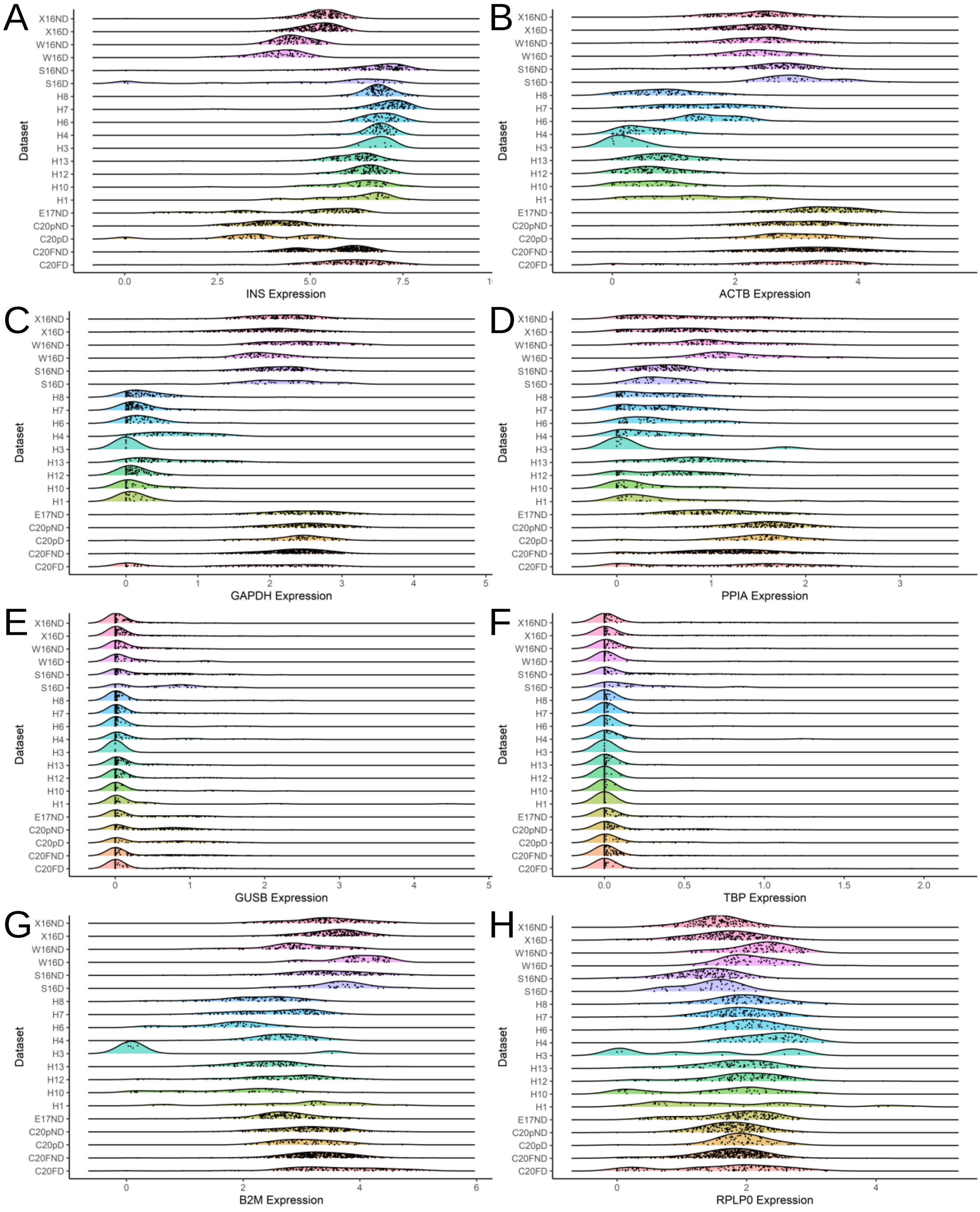


**Supplemental Figure 1: Default normalized INS and housekeeping genes expression by dataset.** (A) INS expression normalized to all genes. (B-H) Housekeeping genes expression normalized to all genes. Gene expression is separated and coloured by datasets. Each point represents one cell. ND = no diabetes, D = type 2 diabetes.
